# Discerning the functional networks behind processing of music and speech through human vocalizations

**DOI:** 10.1101/411074

**Authors:** Arafat Angulo-Perkins, Luis Concha

## Abstract

Musicality refers to specific biological traits that allow us to perceive, generate and enjoy music. These abilities can be studied at different organizational levels (e.g., behavioural, physiological, evolutionary), and all of them reflect that music and speech processing are two different cognitive domains. Previous research has shown evidence of this functional divergence in auditory cortical regions in the superior temporal gyrus (such as the *planum polare*), showing increased activity upon listening to music, as compared to other complex acoustic signals. Here, we examine brain activity underlying vocal music and speech perception, while we compare musicians and non-musicians. We designed a stimulation paradigm using the same voice to produce spoken sentences, hummed melodies, and sung sentences; the same sentences were used in speech and song categories, and the same melodies were used in the musical categories (song and hum). Participants listened to this paradigm while we acquired functional magnetic resonance images (fMRI). Different analyses demonstrated greater involvement of specific auditory and motor regions during music perception, as compared to speech vocalizations. This music sensitive network includes bilateral activation of the *planum polare* and *temporale*, as well as a group of regions lateralized to the right hemisphere that included the supplementary motor area, premotor cortex and the inferior frontal gyrus. Our results show that the simple act of listening to music generates stronger activation of motor regions, possibly preparing us to move following the beat. Vocal musical listening, with and without lyrics, is also accompanied by a higher modulation of specific secondary auditory cortices such as the *planum polare*, confirming its crucial role in music processing independently of previous musical training. This study provides more evidence showing that music perception enhances audio-sensorimotor activity, crucial for clinical approaches exploring music based therapies to improve communicative and motor skills.

## Introduction

HH1uman vocalizations are the most common sound in our environment, and the most ancient expression of music and language^1, 2^. Despite continuous debate regarding the level of functional, anatomical and cognitive independence between speech and music processing^3–7^, it is clear that there are several differences regarding their basic acoustic properties (e.g., temporal, spectral or envelope features) and more importantly, that they do not carry the same type of information^7–15^. These elemental differences support the expression of human musicality defined as the ability or trait that allows us to perceive, create and enjoy music in its various forms^13, 14^. Musical cognitive traits such as melodic, harmonic, timbral or rhythmic processing rely on basic analysis such as relative pitch, beat perception or metrical encoding of rhythm^15, 16^. Interestingly, cortical activity (as measured through blood oxygenation level-dependent [BOLD] signal) is higher in secondary auditory cortices while listening to music as opposed to various types of non-musical human vocalizations—despite these regions being essential for speech processing^17, 18^. The primary auditory cortex (i.e., Heschl’s gyrus), on the other hand, does not show differential activation towards these two types of acoustic categories^19–22^. These results suggest that while music and speech processing share the same basic auditory pathway until the primary auditory cortex, different patterns of activations are observed in other brain regions, with some exhibiting hemisphere lateralization^19–24^.

The immediate question when comparing music and speech refers to vocal music, and whether it also involves different regions when compared to speech, particularly for the case of vocal music with lyrics. Considering voice as the most ancient musical instrument^25–27^, contrasting music and speech processing using vocal sounds has been useful to evaluate brain activity as their spectral and timbre characteristics are pretty similar. This approach has allowed researchers to test both acoustic signals at the same time (e.g., while singing), or to explore them separately (i.e., just linguistic or melodic information) while testing for perception, discrimination, error detection, memory, or different types of production (overt and covert)^10, 24, 28–32^.

Previous findings have shown that regardless of musical stimuli being human vocalizations (such as a syllabic hum or song) there are certain differences in brain activity when compared to that elicited by speech^10, 24, 30, 31^. The next step is to evaluate at which point in the hierarchy of the auditory pathway these two acoustic signals show divergent processing, and if these differences in brain activation are maintained after subtracting some of the basic acoustic properties.

We designed an experimental paradigm including speech sounds, and two different forms of musical vocal sounds, namely one with lyrics (song) and one without (hum). In addition, we included scrambled versions of each category (i.e., speech, song and hum) as a control condition to assess cortical activity related to their temporal structure. To control for semantic content, the stimulation paradigm included 25 different sentences that were sung and spoken by a professional singer. Melodic content was the same for the hummed and sung stimuli.

Finally, considering the literature regarding differences between musicians and non-musicians supporting functional and anatomical differences due to musical training^19, 33–40^, we also explored functional differences associated to musical training and included a group of professional musicians as part of our testing sample. Thirty-three volunteers participated, 16 of whom were professional musicians.

Our main hypothesis was that regions more involved in music processing would show increased activity during both musical categories, in comparison to the speech category regardless of: 1) being produced by the same instrument and 2) the presence or absence of lyrics (i.e., song and hum, respectively). In the same way, we predicted that regions mainly modulated by speech processing would not show differences in their activation in response to similar categories (i.e., song and speech) but would differ with respect to the music category without verbal content (hum). Based on previous results showing that perception of music (in comparison to non-musical human vocalizations) elicited differential brain activity in the *planum polare* and *temporale*, we expected to observe group differences in brain activation patterns for musicians in comparison to non-musicians during vocal-music listening.

## Results

All sound categories produced BOLD signals that were significantly higher than those obtained during the baseline condition (i.e., scanner noise; Figure 1). Analysis of trigger responses to target sounds showed that all subjects responded appropriately (i.e., they were attentive to the stimuli).

**Figure 1.**
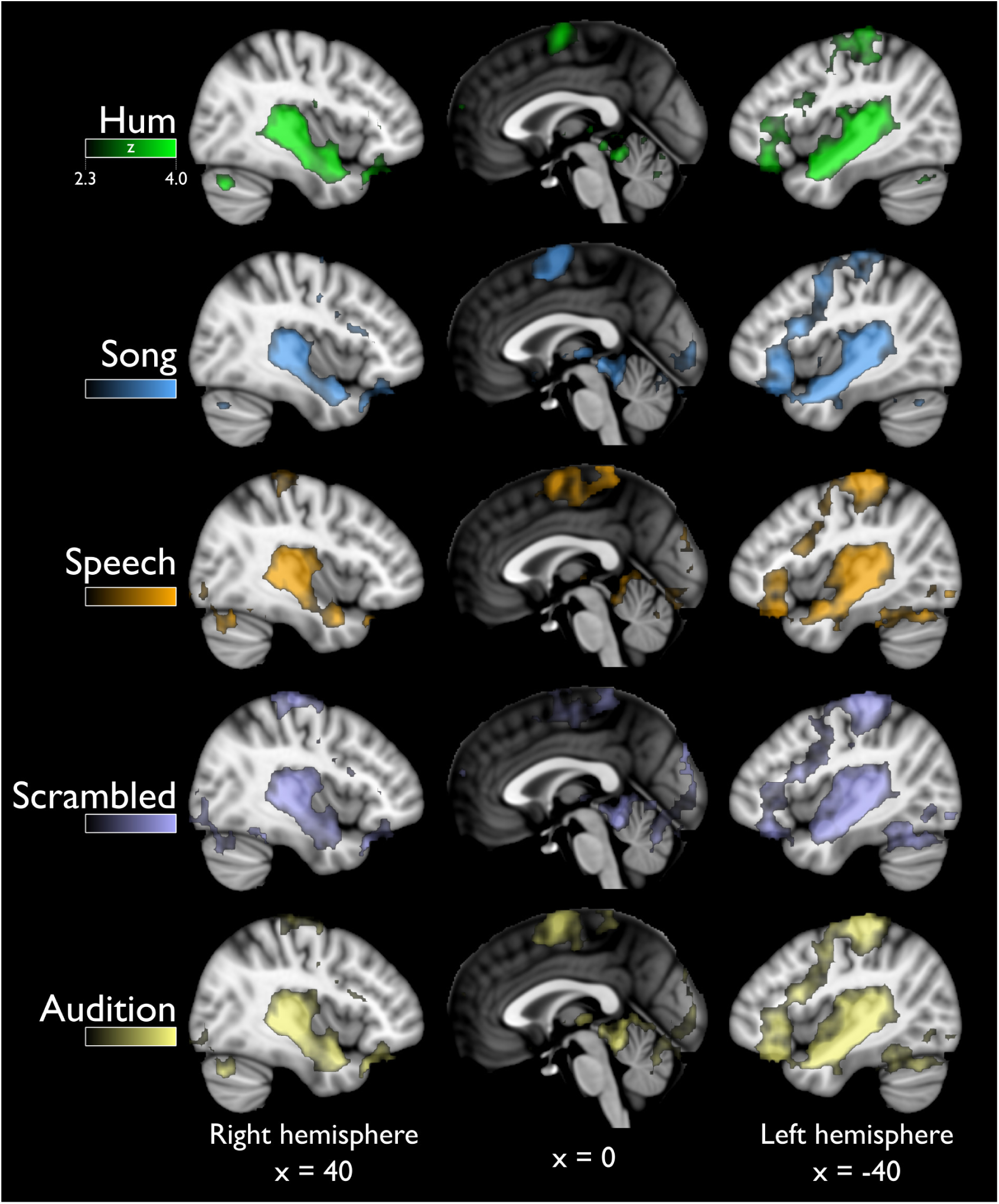
Global activity (above baseline) for all stimuli. Sagittal views showing distributed activation in response to all sound categories, greater than to scanner noise. The last row (Audition) includes all the previous categories. R=right, L=left. Statistical maps are overlaid on the MNI-152 atlas (coordinates shown in mm). For all categories color scale corresponds to z values in the range [2.3 – 4.0], as in top panel.

### Analysis 1: Natural versus scrambled stimuli

All natural stimuli, when compared to the their scrambled counterparts, elicited stronger BOLD activity around (but not in) Heschl’s gyrus in the superior temporal gyrus (STG). However, particular areas were differentially activated by specific categories, as outlined below and visualized in Figure 2. Unthresholded statistical maps for this and all other analyses are available at https://neurovault.org/collections/4116/.

**Figure 2.**
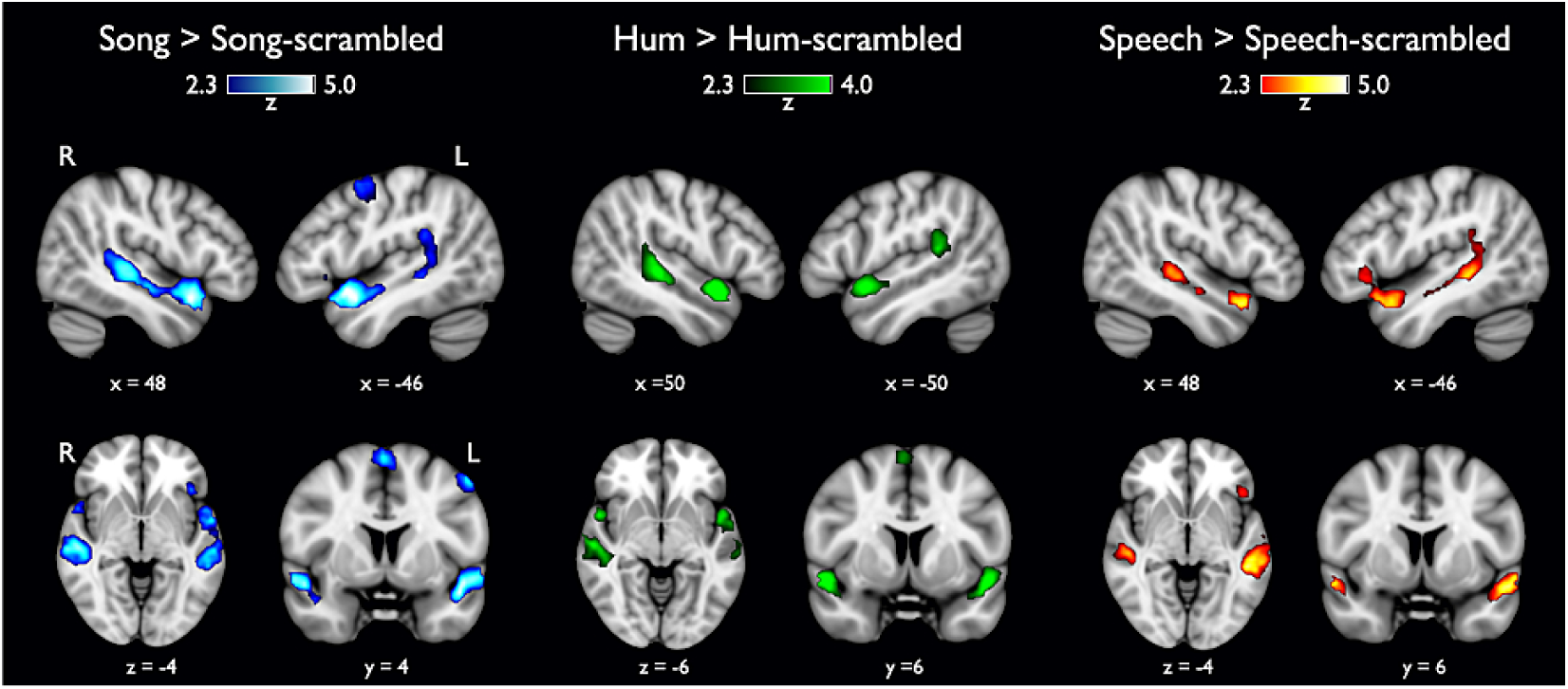
Natural stimuli versus category-specific scrambled stimuli. No differences were found in Heschl’s gyrus; activation in the anterior portion of the superior temporal gyrus, specifically the *planum polare*, was found only in the musical categories.

#### 1.1 Song versus Song-S

The Song > Song-S comparison revealed 4 different clusters, two of them occupying the entire lateral aspect of the STG and the superior temporal sulcus (STS) bilaterally, with the notable exception of Heschl’s gyrus; however, the most significant activation was located within the temporal pole (aSTG; blue color in Figure 2). Cluster volume and coordinates of peak significance for this and all subsequent contrasts are shown in Table 1. The left STG cluster extended into the inferior frontal gyrus (IFG) approximately in Brodmann areas 44 and 45. The third cluster had its most significant voxel in the right supplementary motor cortex (SMA) but it also covered the left counterpart, where it reached the dorsal premotor cortex. The opposite contrast (Song-S > Song) showed 4 clusters covering different regions: bilateral activation of primary auditory regions (Heschl’s gyrus), the insular cortex and two bilateral activations of the middle and inferior occipital gyrus.

**Table 1.** Significant activations for all experiments. Anatomical locations derived from Harvard-Oxford Cortical Structural Atlas. MNI coordinates (*x,y,z*) in mm. Significance threshold: voxel z > 2.3, cluster p < 0.05. Unthresholded statistical maps are available at https://neurovault.org/collections/4116/

| Contrast | Anatomical location | Cluster peak |  |  | Cluster volume<br>(mm<sup>3</sup>) |
| --- | --- | --- | --- | --- | --- |
|  |  | x | y | z |  |
| Song > Song-S | Superior Temporal Gyrus, anterior division \| Temporal Pole (left) | -46 | -10 | -12 | 18112 |
| Song > Song-S | Superior Temporal Gyrus, anterior division \| Temporal Pole (right) | 48 | 12 | -16 | 16280 |
| Song > Song-S | Supplementary motor area (right) | 2 | 2 | 66 | 3616 |
| Song > Song-S | Dorsal premotor cortex (left) | -52 | 2 | 50 | 2648 |
| Song-S > Song | Heschl's gyrus (right) | 56 | -10 | 4 | 10264 |
| Song-S > Song | Heschl's gyrus (left) | -52 | -22 | 14 | 10200 |
| Song-S > Song | Occipital gyrus, middle and inferior division (right) | 48 | -60 | -14 | 7168 |
| Song-S > Song | Occipital gyrus, middle and inferior division (left) | -44 | -58 | -8 | 5032 |
| Hum > Hum-s | Superior Temporal Gyrus, anterior division \| Temporal Pole (right) | 50 | 10 | -14 | 13680 |
| Hum > Hum-s | Superior Temporal Gyrus, posterior division \| *Planum Temporale* (left) | -50 | -38 | 14 | 5664 |
| Hum > Hum-s | Superior Temporal Gyrus, anterior division \| Temporal Pole (left) | -48 | 6 | -12 | 4232 |
| Hum > Hum-s | Supplementary motor cortex (right) | 4 | 4 | 64 | 2512 |
| Hum-s > Hum | Heschl's gyrus (left) | -50 | -18 | 6 | 5936 |
| Hum-s > Hum | Heschl's gyrus (right) | 54 | -10 | 4 | 5376 |
| Speech > Speech-s | Superior Temporal Gyrus, anterior division \| Superior Temporal Sulcus (left) | -54 | 0 | -12 | 17824 |
| Speech > Speech-s | Superior Temporal Gyrus, anterior division \| Superior Temporal Sulcus (right) | 48 | 14 | -20 | 5552 |
| Speech-s > Speech | Heschl's gyrus (left) | 56 | -10 | -4 | 8904 |
| Speech-s > Speech | Heschl's gyrus (right) | -56 | -22 | 8 | 7960 |
| Song > Speech | Superior Temporal Gyrus, posterior division \| Middle Temporal Gyrus, posterior division (right) | 66 | -30 | 2 | 39224 |
| Song > Speech | Temporal Pole \| *Planum Polare* (left) | -48 | 4 | -12 | 15480 |
| Song > Speech | Supplementary Motor Cortex \| Precentral Gyrus (right) | 4 | -2 | 66 | 4048 |
| Speech > Song | Lateral Occipital Cortex, superior division \| Occipital Pole (left) | -22 | -88 | 24 | 25712 |
| Speech > Song | Postcentral Gyrus \| Primary Somatosensory cortex (left) | -20 | -34 | 74 | 13024 |
| Speech > Song | Precentral Gyrus \| Primary motor cortex (right) | 6 | -30 | 60 | 6824 |
| Speech > Song | Heschl's Gyrus \| Parietal Operculum Cortex (right) | 40 | -22 | 14 | 2568 |
| Speech > Song | Insular Cortex (left) | -36 | -8 | 4 | 2248 |
| Hum > Speech | Superior Temporal Gyrus, posterior division \| *Planum Temporale* (right) | 64 | -30 | 10 | 3504 |
| Hum > Speech | Precentral Gyrus \| Premotor cortex (right) | 48 | 0 | 46 | 3048 |
| Hum > Speech | *Planum Polare* \| Temporal Pole (right) | 46 | 4 | -14 | 3008 |
| Speech > Hum | Middle Temporal Gyrus, anterior division \| Temporal Pole (left) | -60 | -10 | -10 | 46888 |
| Speech > Hum | Superior Temporal Gyrus, anterior division \| Superior Temporal Gyrus, posterior division (right) | 62 | -4 | -6 | 19440 |
| Speech > Hum | Postcentral Gyrus \| Primary Motor cortex (right) | 4 | -34 | 60 | 4336 |
| Song > Hum | Superior Temporal Gyrus, anterior division \| Middle Temporal Gyrus, anterior division (left) | -58 | -10 | -10 | 47296 |
| Song > Hum | Superior Temporal Gyrus, anterior division \| Superior Temporal Gyrus, posterior division (right) | 62 | -4 | -6 | 32552 |
| Song > Hum | Inferior Frontal Gyrus, *pars opercularis* \| *pars triangularis* (left) | -56 | 16 | 14 | 3752 |
| Song > Hum | Thalamus (left) | -10 | -16 | 0 | 2624 |

#### 1.2 Hum versus Hum-S

The contrast testing for Hum > Hum-S showed bilateral activation along the STG with three peaks of maximal activation (Figure 2, green colors), two of which corresponded to the left and right temporal pole and the other one to the *planum temporale*. The fourth cluster was located in the supplementary motor area (SMA). No differences were found in Heschl’s gyrus. The inverse contrast testing for Hum-S > Hum evidenced two clusters corresponding to Heschl’s gyrus occupying the adjacent lateral face of the STG.

#### 1.3 Speech versus Speech-S

Functional maps testing for Speech > Speech-S showed two large clusters distributed along the ventral lateral face of the STG reaching the dorsal area of the STS, in the left hemisphere the cluster presented a much larger volume (left 17,824 mm^3^ and right 5,552 mm^3^; Figure 2, warm colors). The left cluster included the IFG, as did the Song > Song-S contrast. The analysis of the opposite comparison (Speech-S > Speech) also revealed two clusters located in the lateral aspect of Heschl’s gyrus (in each hemisphere); cluster locations were almost identical to those found in Hum-S > Hum and in Song-S >Song.

Comparing the original stimuli to a single Scrambled category (as opposed to comparing each natural category with its corresponding scrambled version) showed similar results than those described above, with the exception of the *planum temporale* not showing significantly greater activity for Hum than Scrambled categories (Supplementary Figure S.1).

### Analysis 2: Song and Speech processing

The contrast Song > Speech generated three different clusters. Two clusters were distributed along the STG from the *planum temporale* to the *planum polare* (sparing Heschl’s gyrus); in the right hemisphere this cluster cluster extended to the IFG (Figure 3, cold colors). Peak activation of the third cluster was located over the right supplementary motor area (SMA), however it also covered part of the right premotor cortex (PMC). The opposite contrast (Speech > Song; Figure 3, warm colors) elicited activation of the superior parietal lobe corresponding to the primary somatosensory (S1) cortex, extending slightly into the primary motor cortex (M1). We found another cluster located in the lateral occipital cortex of the left hemisphere covering the primary visual cortex (V1); these areas were also active in the right hemisphere as an extension of the right parietal cluster. A group of voxels were found bilaterally in the posterior portion of the Heschl’s gyrus, as extensions of clusters in the insular cortices.

**Figure 3.**
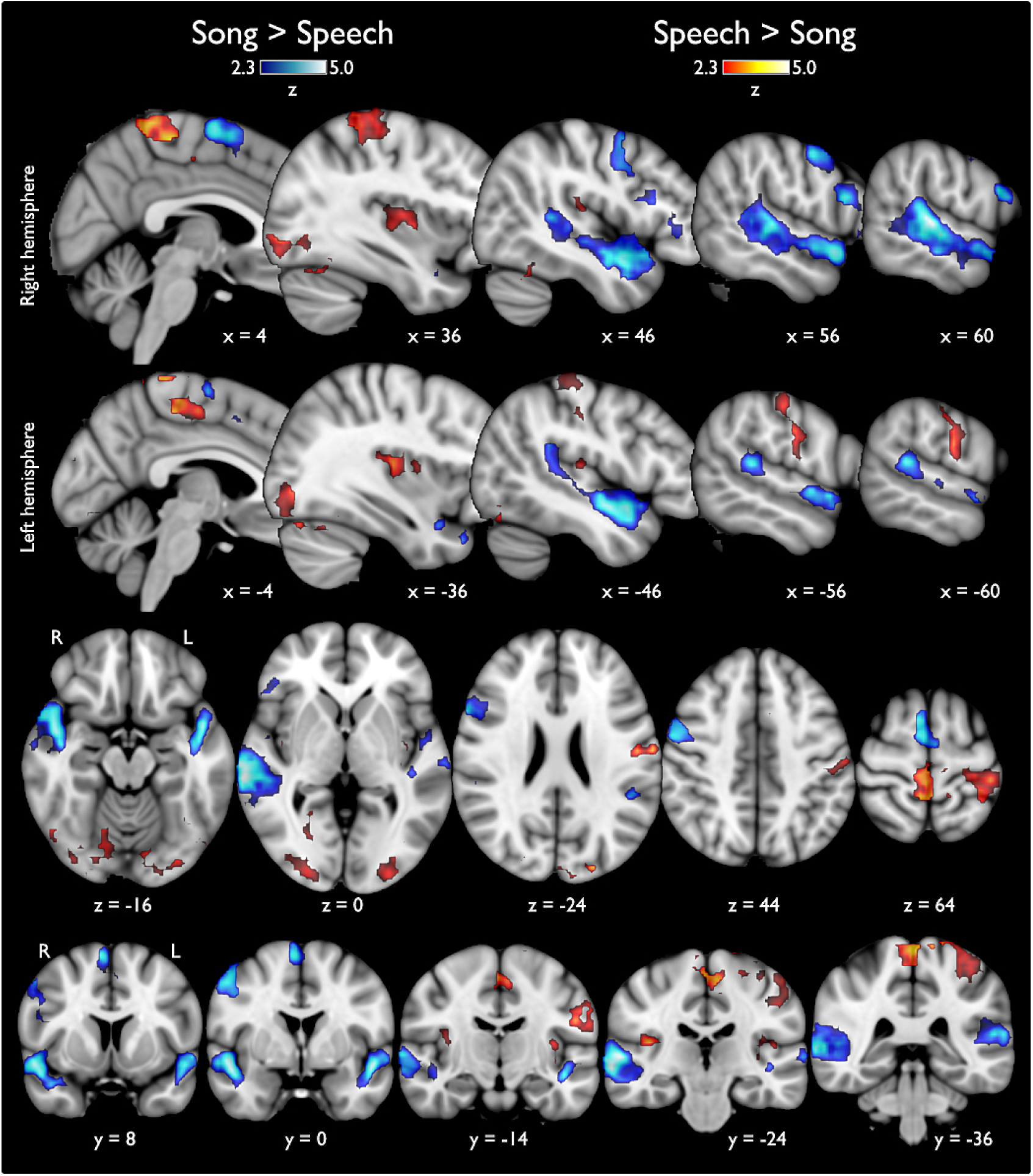
Statistical activation maps for differences between vocal music and speech sounds. Blue colors indicate those regions that were more active while listening to song stimuli as compared to speech (i.e. Song > Speech); warm colors show the opposite comparison (Speech > Song). Top and middle panels show lateral progression in the sagittal plane, from medial to lateral regions in each hemisphere. Axial and coronal views are presented in the inferior two panels to facilitate visual inspection of all clusters).

### Analysis 3: Hum versus speech

Functional maps testing for Hum > Speech (Figure 4; top panel, green colors) revealed activation along the right STG including the *planum polare* and *planum temporale*. We also found activation in the precentral gyrus, specifically in the PMC. Using an uncorrected threshold p < 0.001 (outlined in bright green in Figure 4), we observed bilateral activation of the *planum polare* and right SMA. The opposite comparison (Speech > Hum) revealed three clusters. Two were distributed in different areas of the STG including bilateral activation of Heschl’s gyrus, the lateral aspect of the STG, the STS and MTG (Figure 4, top panel, warm colors). The cluster in the left temporal lobe partially extended to the IFG. The other cluster was distributed mainly in the left postcentral gyrus (covering S1) and the ventral division of M1.

**Figure 4.**
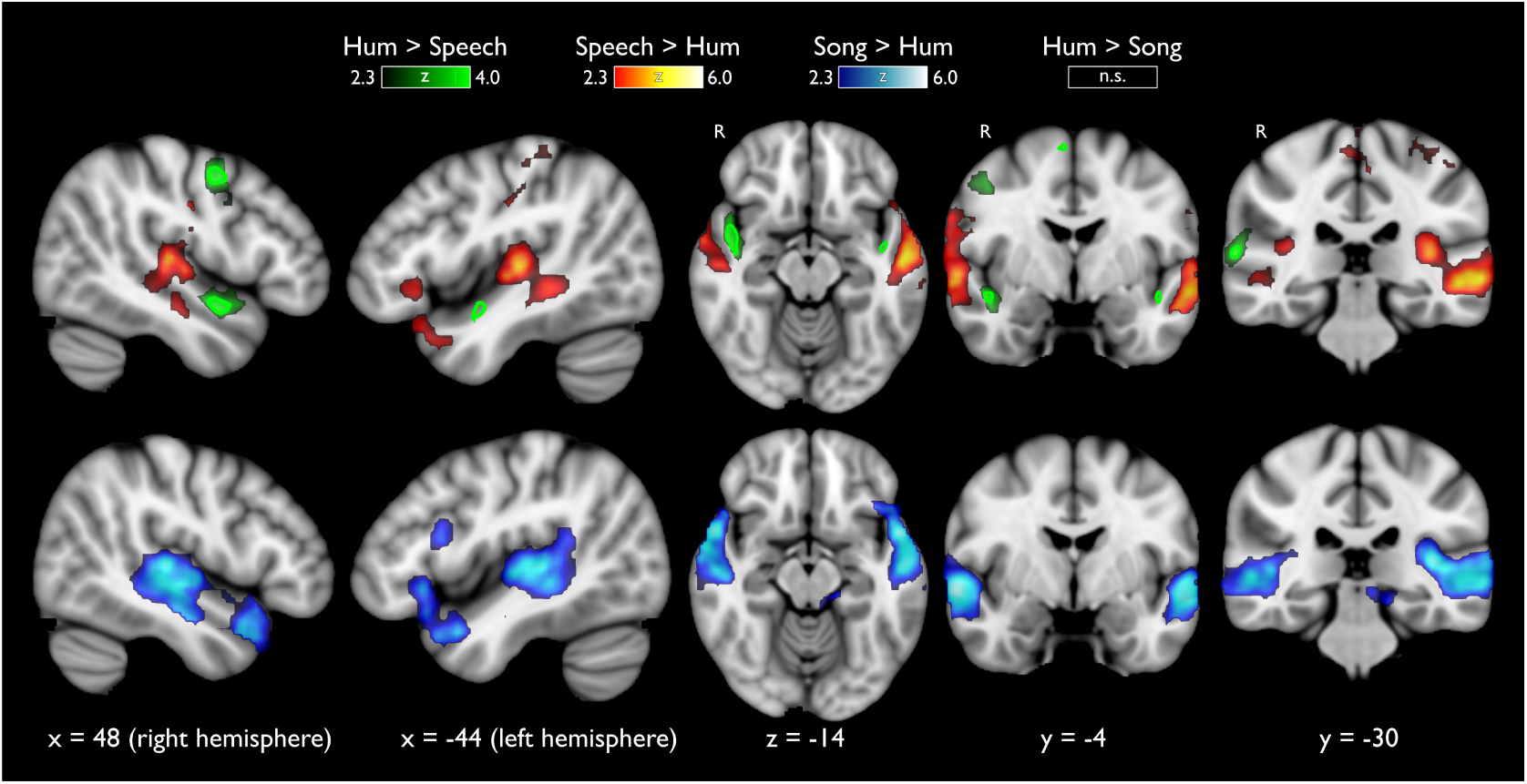
Top panel: Hum versus Speech. Listening to hummed songs recruited a right hemisphere network (green colors) including the right *planum polare*, and PMC, as compared to listening to speech stimuli. Uncorrected results (p < 0.001; shown in bright green) show bilateral activation of the *planum polare* and activation of SMA. The opposite contrast (i.e., Speech > Hum; warm colors) shows activation in the left hemisphere including the IFG, *planum temporale*, S1 and bilateral activation of Heschl’s gyrus and STS. Bottom panel: Vocal-music with and without lyrics. Lateral, axial and coronal views showing in blue the cortical regions modulated preferentially to Song as compared to Hum stimuli, extending over temporal and frontal cortices. The opposite contrast (Hum > Song) did not show any significant activity.

### Analysis 4: Comparing vocal-music with and without lyrics

The results obtained from the contrast Song > Hum (Figure 4; bottom panel, cold colors) generated four clusters, two of them covered anterior and posterior regions of the STG (including Heschl’s gyrus was included in both hemispheres). These activations reached the STS and part of the dorsal portion of the MTG. In the left hemisphere we also found activation in the IFG (*pars opercularis* and *triangularis*) and in the thalamus. The reverse contrast (Hum > Song) showed no significant activations.

### Analysis 5: Influence of musical training

All the previously described contrasts were used to test for differences between musicians and non musicians. We did not find any statistically significant differences between groups (see Methods for details).

### Analysis 6: Functional characterization of the superior temporal gyrus

To further analyze BOLD signal changes derived from the different sound categories along the auditory cortices we used an unbiased approach to compare BOLD changes in the *planum polare, planum temporale* and Heschl’s gyrus. Figure 5 illustrates the percentage of BOLD signal change for each category. We found statistically significant differences between Song and Speech in the left and right *planum polare*, and also between Song and Hum; the lowest levels of activation within the *planum polare* were observed in the Speech category. BOLD responses in Heschl’s gyrus (A1) were lower in response to Hum as compared to those elicited by Song or Speech, bilaterally; there were no differences in A1 activation between the last two categories. In parallel with this observation, acoustic feature analyses showed that Hum differs substantially from Song/Speech in several aspects, such as zero-cross, and spectral spread, brightness, centroid, kurtosis and flatness (see Supplementary Figure S.2).

**Figure 5.**
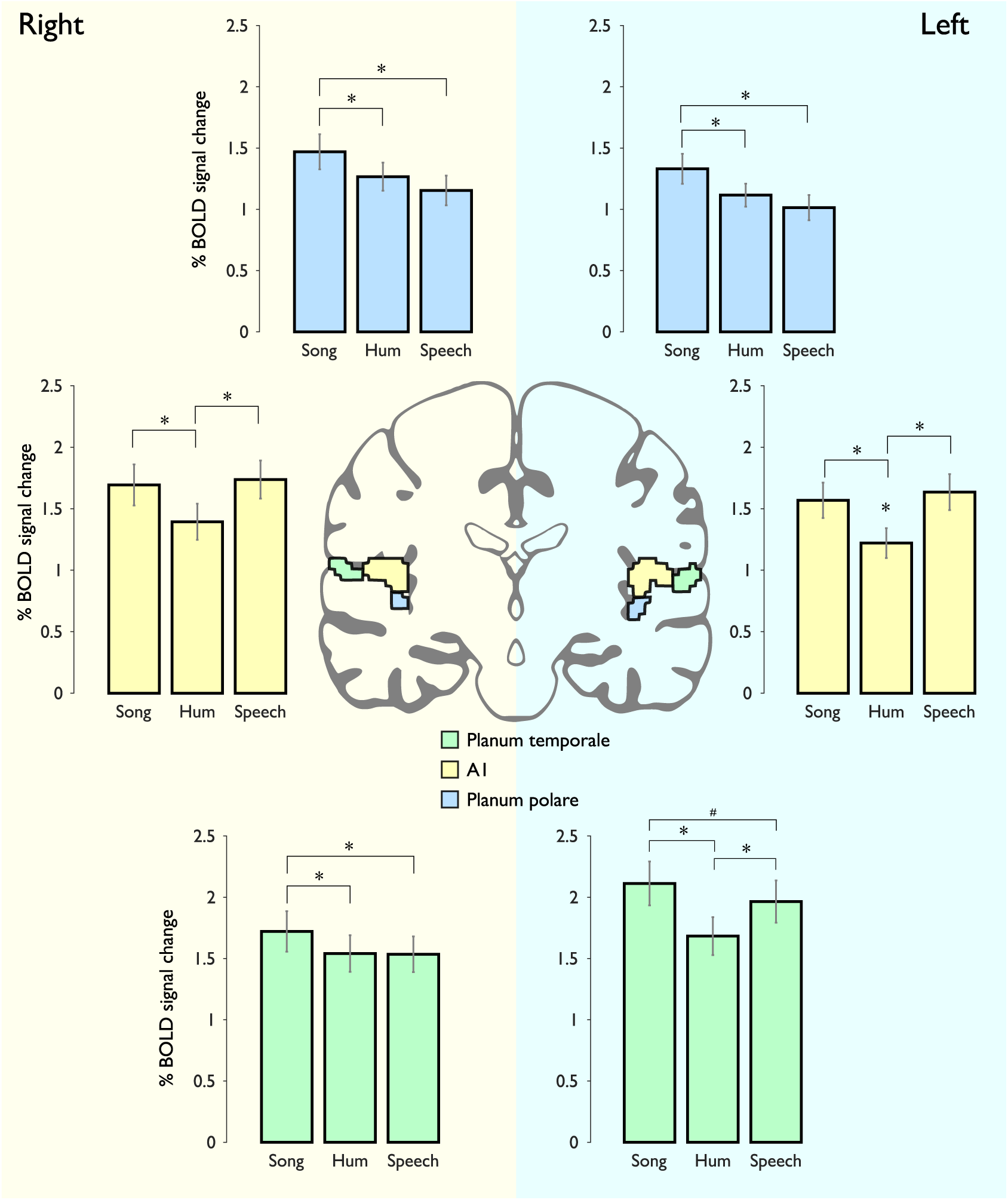
ROIs of Heschl’s gyrus, *planum polare* and *planum temporale*. Right and left hemispheres are shaded in different colors. Top panel: The percentage of BOLD signal modulation in the *planum polare* (blue color) showed similar patterns between the right and left hemisphere, the highest level of activation in this region corresponded to Song, followed by Hum and Speech sounds; statistical differences were found between Song and Speech. Middle panel: BOLD signal changes in the right and left Heschl’s gyrus (yellow color), no differences between Song and Speech stimuli. Bottom panel: The left and right *planum temporale* (green color) exhibited different patterns of activation, in the right hemisphere Song stimuli elicited the highest values, while in the left hemisphere there were no differences between them. * indicates significant difference from the rest of the stimuli (p<0.0028; Bonferroni correction); # stands for uncorrected p<0.05.

The *planum temporale* showed different patterns between hemispheres. The right *planum temporale* was equally active during Hum and Speech conditions in contrast to Song; however, in the left *planum temporale* we observed a similar pattern to that seen in A1, namely it was preferentially activated by Song and Speech categories (Figure 5; lower panel). From all comparisons among the three different regions in the STG, the highest activations were found in the left *planum temporale*.

## Discussion

In this study we aimed to further explore the neural basis of music and speech perception evaluating whether the functional selectivity would exhibit a similar pattern when comparing vocal music, with and without lyrics, versus speech. On the basis of previous observations and other studies using similar approaches^10, 19, 24, 28–31^, we designed a passive listening stimulation paradigm that included different vocal sound categories (i.e., Song, Hum and Speech), all of them compared against their respective scrambled version. This strategy allowed us to control for melodic and semantic content by using the same sentences in Speech and Song categories, and the same musical excerpts for Hum and Song stimuli, as well as partially controlling for differences of timbre (using the same voice to produce all the stimuli). Notably, we eliminated some of the basic attributes related to the temporal structure of the stimuli by subtracting the brain activity from all the scrambled versions, an aspect that had not been previously evaluated.

We compared the original acoustic stimuli with their scrambled versions (e.g., Song > Song-S) to evaluate the effect of temporal structure on cortical activity, while controlling for basic acoustic features. Interestingly, we found no differences in Heschl’s gyrus activation, whereas we found bilateral activation of secondary auditory cortices (i.e., *planum polare* and *temporale*), IFG, SMA and dorsal PMC. This result confirms that while basic aspects of sound decomposition are performed in primary auditory cortex, processing of temporally-complex sounds, such as vocal music and speech, requires cortical regions beyond Heschl’s gyrus^20, 28^. Activation maps resulting from comparing the natural and scrambled versions of each category showed that while all types of stimuli recruited the STG, only musical stimuli involved activation of SMA and PMC. By comparing the statistical maps derived from the contrasts Song > Song-S and the contrast testing Speech > Speech-S, we found four distinct regions that were more active in the Song contrast, corresponding to the *planum polare* in both hemispheres, and the left PMC and SMA. These results were consistent with previous studies reporting BOLD signal increases in these brain areas in response to temporal, semantic, syntactic and melodic-contour manipulations of musical stimuli^23, 41, 42^.

The opposite contrasts comparing the scrambled versions (e.g., Song-S > Song) revealed consistent higher BOLD modulations in Heschl’s gyrus for all the three types of vocal stimuli. It is well known that the primary auditory cortex presents a tonotopical organization sensitive to frequency tuning and spectro-temporal modulations^43–46^. However, it also contributes to the construction of auditory objects, and it has therefore been suggested that less meaningful sounds require more neural resources to extract information from them, in an attempt to elaborate predictions about their possible identity as an auditory object^47, 48^.

Analyses 2-5 query brain regions involved in high-level acoustic processes (e.g., abstract representations). The *planum polare* and *planum temporale* maintained their selectivity for vocal music when we compared them to Speech. This result corroborates that these auditory regions are particularly relevant for music processing independently if the music is produced by the human voice or other instruments such as piano or violin^19–21, 24^. However, each region presented a different response to the music stimuli: while the *planum polare* in both hemispheres exhibited higher levels of activation for both types of musical stimuli, the left *planum temporale* shows equal activations elicited by Speech and Song categories, This suggests a preference of the left *planum temporale* for stimuli with verbal content (whether musical or not), possibly associated with representations of lexical or semantic structures^49–51^. Moreover, the highest activation of the right *planum temporale* was elicited by Song stimuli, (whereas no differences were found between Hum and Speech), which can be attributed to a summatory effect. This evidence supports a possible functional specialization within the right *planum temporale* as has been suggested in other studies, but also a role in higher order perceptual analysis^28–30, 51^.

Previous studies have reported activation in the *planum polare* during evaluation of vocal music tasks, overt and covert production of songs and sentences and melodic repetition and harmonization in musicians^10, 30–32^. In line with those reports, our results suggest that the *planum polare* might be playing an intermediate role between the primary auditory cortex and other associative cortices, possibly extracting information (such as melodic patterns or pitch-interval ratios) required for further processing leading to perceptual evaluations (e.g., same-different task), vocal production, and sensory-motor coordination to reproduce melodic or rhythmic sounds^30, 31^.

Voice perception elicits activity of specific motor cortices involved in speech production^52^. Our results extend previous observations, as we show that motor cortex activity is greater when the voice heard produces musical patterns, independently of whether the music was only hummed or sung with lyrics and also regardless of musical training. This was particularly evident in cortical motor regions such as the SMA and the PMC, which are related to beat and rhythm perception^53–58^. Considering that we gave explicit instructions to avoid movement during the scanning session, the increased activation of this audio-motor regions could be generated by the perception of the rhythmic patterns to enhance pulse and meter perception abilities^53–55, 57, 58^.

Activation of the right IFG in both musical conditions confirms previous studies suggesting this region as a rightward homologue of the left IFG, processing dynamic pitch contour possibly based on the temporal coherence of rhythmic and melodic variations^59–61^. This result is in line with the differential role of the IFG suggested for pitch processing during speech and vocal music listening (left-speech, right-song)^24^.

In a previous study using different types of instrumental music we demonstrated that, as compared to non-musicians, individuals with musical training show greater activity of the *planum polare* bilaterally, and of the right *planum temporale* when listening to musical stimuli^19^. In this study, however, we did not find differences related to musical expertise in any of the analyses using vocal music. This apparent discrepancy may be explained by: 1) the relatively smaller size of our current sample; 2) the diversity of musical training in the group of musicians (Supplementary Table S.1), which may induce functional plastic changes related to instrument-specific tuning^39, 40^; and 3) common vocal training. Most individuals (musicians or not) sing in different everyday situations since very early ages (e.g., birthdays, hymns, cheers). Learning to play an instrument is, in contrast, not a universal activity. Therefore, we can not draw definite conclusions regarding this particular comparison. To do so, musicianship should be assessed more thoroughly, and this type of study should be performed on a group of professional singers.

Speech processing involves many cortical regions (e.g., primary auditory cortices, left STG, *planum temporale*, STS, IFG, postcentral gyrus and the ventral division of M1)^62, 63^, which are recapitulated in our results. Interestingly, we found bilateral activation of the Heschl’s gyrus when we compared Speech sounds with Song and Hum categories after subtracting the basic acoustic attributes. This partially confirms that Heschl’s gyrus is involved in complex acoustic analyses specifically related to speech processing, such as phoneme perception^10, 32, 64^.

Speech stimuli also induced greater activity within S1, as seen in the comparison Speech > Hum, which was not observed when the speech stimuli were musically modulated (i.e., Song > Hum). In conjunction with the increased activity seen in M1, these results hint at an enhanced sensorimotor feedback, probably facilitating motor patterns normally required for phonation^65, 66^.

Many of our results are consistent with those described by Schön et al.^10^. Our work confirms and extends such study, yet their results are particularly informative about the neurobiological substrates involved in song, vocalise and speech processing specifically during a cognitive task (i.e., evaluating similarities and differences among two auditory stimuli). Through the task-free nature of our design, we found more involvement of the posterior portion of the superior temporal gyrus (including Wernicke’s area) during listening of vocal music. Similarly, said study did not show different cortical activity for the Speech > vocalise comparison. The equivalent condition in our study (i.e., Speech vs Hum) resulted in a distributed pattern including several regions such as the left inferior frontal gyrus, the superior precentral gyrus, inferior pre-post central sulcus, anterior superior temporal gyrus, both Heschl’s gyri, among other areas. This difference could be related to their experimental condition using syllabic singing (i.e., vocalise *-vi vi vi-*), which is distinct to humming, not only because the vocalization includes proper syllables (which are part of words), but also for muscle coordination patterns (e.g., open versus close mouth) and other basic acoustic properties (e.g., humming is more monotonic with slight tone variations).

An independent and very recent study compared cortical activity in response to instrumental music, speech and *a capella* singing, and also showed that the *planum polare* was preferentially activated when subjects listened to either kind of musical stimuli (sung or instrumental)^67^. As described above, the cortical areas that we found to be most active in response to listening to music include not only the temporal lobe, but also motor and pre-motor regions. The fine-grained matching of our stimuli (all performed by the same singer) may be attributable, in part, to our more extended findings.

### Limitations of the Study

Together, these data demonstrate different cortical regions that are preferentially modulated by particular sounds, whether they are music, speech or a mixture of the two. However, the anatomical resolution given by the technique does not allow us to distinguish finer anatomical details regarding the specific distribution of the statistical maps. For the same reason, we cannot elaborate a more detailed analysis regarding the participation of the primary auditory cortex in Speech and Song conditions, for example. Temporal resolution is also limited in all fMRI studies, and there is a wealth of information from rapid temporal fluctuations present in music and language that are not easily addressed with this technique. As such, our results can only reveal relatively long-term changes of brain activity in response to listening to specific sound categories, and our conclusions will benefit greatly from other methods with high temporal resolution, such as (magneto-) electrophysiological recordings. Finally, although stimuli of all three categories were produced by the same singer, we acknowledge that other acoustic features were not homogenized to control for all acoustic parameters that could potentially differ between categories. While some differences in cortical activity may be explained by these low-level features, our main findings are better explained by higher-level, time-varying acoustic features that are characteristic of each category.

### Conclusions

We demonstrated distributed brain regions for the processing of vocal music with higher activations in the right hemisphere, involving frontal and temporal cortical areas involved in auditory and motor processing. Our data also corroborated music selectivity, particularly in the anterior portion of the secondary auditory cortex (*planum polare*), which was independent of whether the musical stimuli contained lyrics or not. Finally, these results provide more evidence supporting music-based therapies as a complementary way to address motor impairments^68–71^ and communicative skills, as has been demonstrated in social behaviors in children with autism^72–74^.

## Methods

### Participants

Thirty-three healthy, right-handed volunteers, age 28±8 years (range: 20 to 42 years; 17 women), participated in this study. Seventeen volunteers were non-musicians (age 27±6 years; range: 20-45 years; 9 women), who had not received extra-curricular music education beyond mandatory courses in school. Musicians (16 volunteers; age 28±7 years; range: 20-42 years; 8 women) had received at least 3 years of formal training/studies in music (either instruments or singing) and were currently involved in musical activities on a daily basis (Supplementary Table S.1). Groups did not differ in terms of age or gender. All volunteers were native Spanish speakers, self-reported normal hearing (which was confirmed during an audio test within the scanner), were free of contraindications for MRI scanning and gave written informed consent before the scanning session. The research protocol had approval from the Ethics Committee of the Institute of Neurobiology at the Universidad Nacional Autónoma de México and was conducted in accordance with the international standards of the Declaration of Helsinki of 1964.

### Experimental design

The vocal music paradigm used a pseudo-randomized block design; each block lasted 15 seconds and included 5-7 stimuli from the same category (Figure 6, panel A). Six different categories were included: Hum, Song, Speech, and their scrambled counterparts (5 blocks each along the paradigm). Additionally, 5 blocks of silence (each lasting 15 seconds) were interspersed throughout the stimulation paradigm. The stimulation protocol had a total duration of 10 minutes.

**Figure 6.**
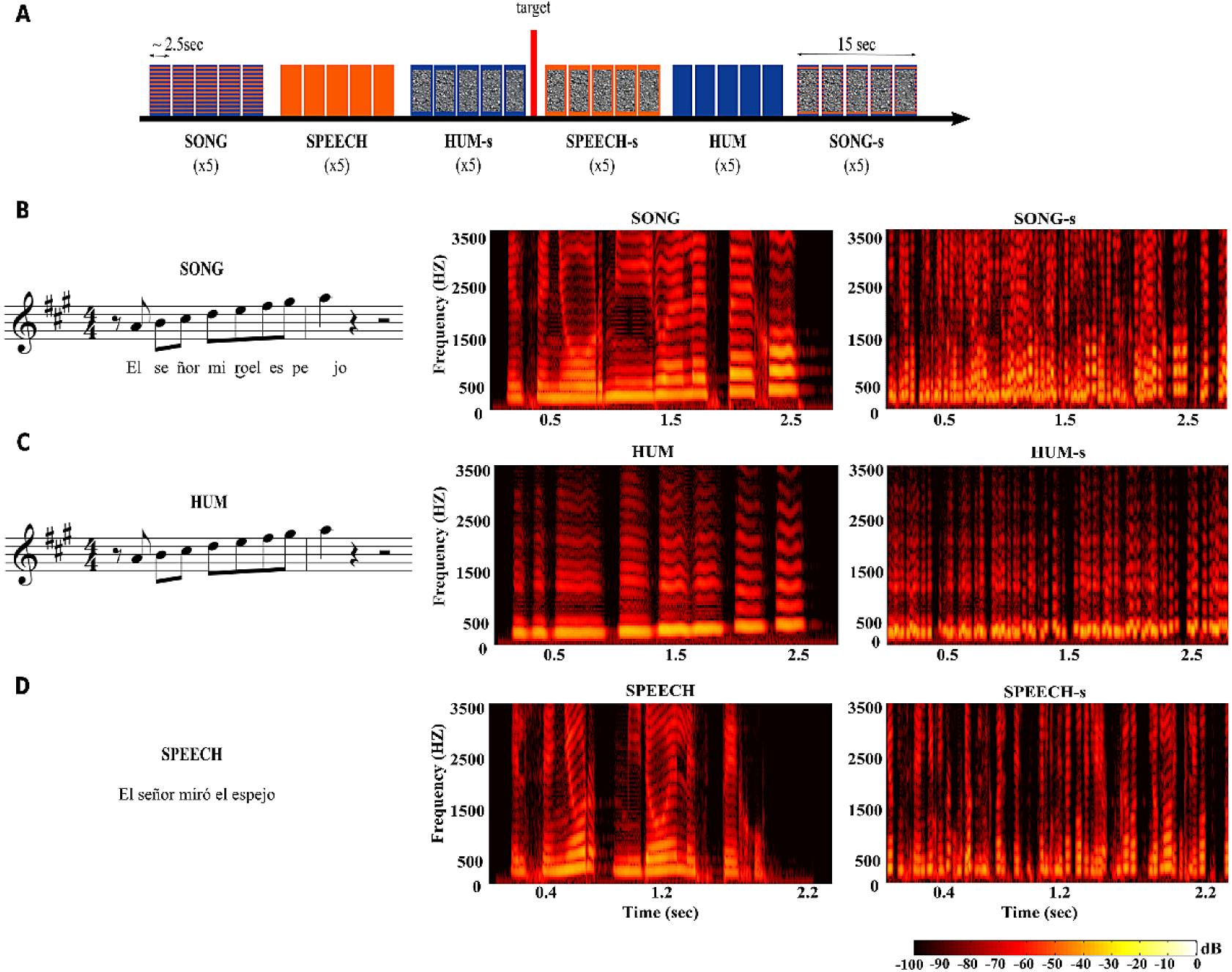
(A) Auditory stimulation paradigm. Each block (∼15 sec) included 5-7 different stimuli from the same category. The paradigm included 5 blocks of each of the 6 sound categories, for a total duration of 10 min. Blue indicates musical sounds, orange for speech stimuli, and scrambled gray indicates the scrambled version of each category (e.g., Hum-S). The target sound was presented between two randomly selected blocks on five occasions throughout the paradigm. (B-D) Example stimulus in three versions: Song (B), Hum (C) and Speech (D). All stimuli were produced by the same singer. Spectrograms show the frequency structure (y-axis) over time (x-axis), with colors representing the relative amplitude of each frequency band. Left column shows natural categories; right column shows the temporally scrambled condition. “*El señor miró el espejo*”: “The man looked at the mirror”.

The stimulation paradigm was presented with E-prime Study Software (version 2.0; Psychology Software Tools, Sharpsburg, PA) binaurally through MRI-compatible headphones (Nordic NeuroLab, Bergen, Norway) that attenuated acoustic interference (∼20 dB) generated by the gradients. While not used as a formal test for normal hearing, we performed a short audio test (1 min) inside the scanner, using similar stimuli, to evaluate whether volunteers could hear and recognize the different sounds inside the scanner; the volume was deemed comfortable but sufficiently high to mask the noise generated by the imaging acquisitions. Subjects were instructed to pay attention to the stimuli and to press a button with the index finger of their right hand every time they heard a pure tone (500 Hz, 500 ms duration), which was presented 5 times randomly throughout the paradigm. We used this strategy to ensure attention throughout the stimulation paradigm, and we used a pure tone as it is clearly different from the rest of our stimuli and therefore easily recognizable. Subjects kept their eyes open during scanning.

#### Acoustic stimuli

Vocal-music paradigm (Figure 6, panel A): Auditory stimuli consisted of short excerpts of 2.8±0.5 seconds, normalized to −30 dB using Adobe Audition (Adobe Systems). MIRToolbox (version 1.7)^75^, implemented in MATLAB (Mathworks, Natick, MA), was used for acoustic analyses.

All stimuli (divided into Song, Speech and Hum categories; Figure 6, panel B-D) were produced by a professional female singer after a period of training to avoid any emotional emphasis during production (e.g., affective prosody or emotional intonation). The sentences used in the Song and Speech categories were novel and carefully selected from a pool of 80 phrases in a pilot test, where 30 listeners (15 women), who did not participate in the main experiment, rated the emotional valence (e.g., from very emotional to neutral) and the complexity of each stimulus. Finally, we selected for use in the imaging experiments the 35 sentences considered the most neutral and simple, both in their grammatical structure and meaning (e.g., *“La alfombra está en la sala”* - “The rug is in the living room”).

#### Hum

This category included 25 novel musical sequences that we had previously used^19, 76^; all melodies followed rules of Western tonal music. The singer hummed these melodies with her mouth closed (i.e., no syllable was used). Each block in this category consisted of five different melodies.

#### Speech

35 Spanish sentences were included in this category. Given the slightly shorter duration of spoken sentences as compared to their sung versions, seven phrases were presented per block.

#### Song

The same 25 musical sequences from the hum category were used as melodies to produce the sung versions of 25 sentences used in the speech condition. Five songs were included per block.

#### Scrambled stimuli

(Figure 6, panel B-D): The scrambled versions of each stimulus served as a control condition for the three main categories. Small fragments (50 ms) with 50% overlap were randomly repositioned temporally within an interval of one second using a freely available Matlab toolbox (http://www.ee.columbia.edu/∼dpwe/resources/matlab/scramble/). This procedure retained low level acoustical attributes (i.e., pitch, duration, loudness, and timbre) but rendered the stimuli unintelligible by disrupting their temporal organization (i.e., melody and rhythm) and therefore, their high-level perceptual and cognitive properties. The scrambled counterparts of each of the original sound categories are identified as: 4) Hum-S, 5) Speech-S and 6) Song-S.

### Image acquisition

All images were acquired at the National Laboratory for Magnetic Resonance Imaging using a 3T Discovery MR750 scanner (General Electric, Waukesha, Wisconsin) with a 32-channel coil. Functional volumes consisted of 50 slices (3 mm thick), acquired with a gradient-echo, echo-planar imaging sequence with the following parameters: field of view (FOV) = 256 ×256 mm^2^, matrix size = 128 ×128 (voxel size = 2 × 2 ×3 mm^3^), TR = 3000 ms, TE = 40 ms. To improve image registration we also acquired a 3D T1-weighted volume with the following characteristics: voxel size = 1 ×1 ×1 mm^3^, TR=2.3 s, TE=3 ms. The total duration of the experiment was 18 minutes. All imaging data is available upon request and will be made freely accessible upon acceptance at openneuro.org.

### Image processing and statistical analyses

Anatomical and functional images were preprocessed using *fsl* tools (version 5.0.9, fMRIB, Oxford UK). Images were corrected for movement and smoothed using a 5-mm FWHM Gaussian kernel; spatial normalization was performed using the MNI-152 standard template as reference. fMRI data analysis was conducted using FEAT (FMRI Expert Analysis Tool) version 6.00. Statistical analysis was performed using the general linear model. For each subject (first level analysis) the three original acoustic categories were modelled as explanatory variables (EV), along with their three scrambled counterparts (Analysis 1) or a single EV comprising the three scrambled categories (Scrambled; Analyses 2-6). Using a single scrambled category in the statistical model allows for the direct comparison of the natural categories controlling for variance explained by low-level acoustical features (e.g., [Hum>Scrambled] > [Song>Scrambled] is equal to Hum>Song). Target stimuli were included as a nuisance regressor. Statistical maps of between-category differences for each subject were generated using a fixed-effects model, and the resulting contrasts were entered in a random-effects model for between-subject analyses using FLAME^77, 78^; musical expertise was included as a group factor at this level. We used random field theory^79^ to correct for multiple comparisons (voxel z > 2.3, cluster p < 0.05) unless otherwise specified.

## Analyses

### Analysis 1 (natural versus scrambled stimuli)

By disrupting the global perception of the stimuli through temporal scrambling, while leaving low-level acoustic features untouched, we searched for brain areas that showed greater activation by natural stimuli compared to their scrambled counterparts in each category. The contrasts included were: 1.1 Song vs. Song-S; 1.2 Hum vs. Hum-S and 1.3 Speech vs. Speech-S.

### Analysis 2 (Song versus Speech)

We searched for differences in brain activity in response to listening to song and speech stimuli, both of which include semantic information but differ in temporal (e.g. rhythm) and spectral (e.g. pitch modulation) content. The contrasts included were: 2.1 Song vs. Speech; 2.2 Speech vs. Song.

### Analysis 3 (Hum versus Speech)

These two categories differ in terms of temporal, spectral and semantic content. The contrasts included were: 3.1 Hum vs. Speech; 3.2 Speech vs. Hum.

### Analysis 4 (Hum versus Song)

This analysis aimed to find differences in brain activity in response to listening to two categories of melodic sounds that differed in semantic content. The contrasts included were: 4.1 Hum vs. Song; 4.2 Song vs. Hum.

### Analysis 5 (musicians versus non-musicians)

We evaluated differences between groups to identify whether musical training changes the patterns of activation during the perception of different types of human vocalizations. We included the following comparisons: 5.1 Song vs. Speech | musicians > non-musicians; 5.2 Speech vs. Song | musicians > non-musicians; 5.3 Hum vs. Speech | musicians > non-musicians; 5.4 Speech vs. Hum | musicians > non-musicians: 5.5 Song vs. Hum | musicians > non-musicians; 5.6 Song vs. Hum | musicians > non-musicians.

### Analysis 6 (Functional characterization of the Superior Temporal Gyrus)

Finally, we also performed an independent functional ROI (region of interest) analysis of the *planum polare, planum temporale* and Heschl’s gyrus. ROIs were derived from the Harvard-Oxford Probabilistic Anatomical Atlas thresholded at 33%; statistical significance threshold was set at p< 0.0028, considering eighteen comparisons were performed (i.e., three categories in six ROIs). This analysis explored BOLD signal modulations to melodic and non-melodic vocal sounds in three different auditory regions.

## Acknowledgements

This study was sponsored by Conacyt (IE252-120295 and 181508) and UNAM-DGAPA (I1202811 and IN212811). We thank the Posgrado en Ciencias Biomédicas of the Universidad Nacional Autónoma de México (UNAM) and CONACYT for Graduate Fellowship 233109 to A.P.A. We are grateful to all the volunteers who made this study possible. We also thank Pézel Flores for helping us to select, evaluate and record all the vocal stimuli; Daniel Ramírez Peña for his valuable help with the validation and preprocessing of all the acoustic stimuli; Edgar Morales Ramírez for his permanent technical assistance; Leopoldo González-Santos, Erick Pasaye, Juan Ortíz-Retana and the support team from the Magnetic Resonance Unit (UNAM, Juriquilla), for technical assistance; and Dorothy Pless for proof-reading and editing our manuscript. Finally, we thank the authorities of the Music Conservatory “José Guadalupe Velázquez”, the Querétaro School of Violin Making, and the Querétaro Fine Arts Institute for their help with the recruitment of musicians. The authors declare no competing financial interests.

## Author contributions statement

A.A-P and L.C conceived the experiments and analysed the results, A.A-P conducted the experiments. Both authors reviewed the manuscript.

## Supplementary Material

### Data availability

- Raw fMRI data is available upon request, and will be made freely available for download from openneuro.org upon acceptance as a peer-reviewed article.
- Unthresholded statistical maps for all contrasts described in the manuscript are available at https://neurovault.org/collections/4116/

**Table S.1.** Musical training information

| Subject | Practice (y) | Age of onset (y) | Weekly practice (h) | Weekly listening (h) | Instruments played | Main expertise |
| --- | --- | --- | --- | --- | --- | --- |
| 1 | 13 | 7 | 5 | 14 | (5) guitar, keyboard, bass, harmonica and flute | (1) guitar |
| 2 | 30 | 12 | 10 | 30 | (3) flute, guitar, piano and violin | (1) violin |
| 3 | 8 | 12 | 5 | 10 | (7) piano, guitar, bass, ukulele, saxophone, synthesizer and percussions | (6) piano, guitar, bass, ukulele, saxophone and synthesizer |
| 4 | 13 | 7 | 9 | >30 | (3) piano, psaltery, voice | (3) piano, psaltery, voice |
| 5 | 8 | 18 | 15 | 50 | (2) guitar and saxophone | (1) saxophone |
| 6 | 12 | 12 | 12 | 14-20 | (4) guitar, bass, mandoline and keyboard | (1) guitar |
| 7 | 22 | 5 | 5 | 20 | (4) piano, guitar, bass, harmonica | (2) guitar and piano |
| 8 | 27 | 13 | 8 | 40 | (2) guitar and violin | (1) guitar |
| 9 | 21 | 6 | 15 | 50 | (4) piano, guitar, bass, percussions | (2) piano and bass |
| 10 | 13 | 8 | 32 | 50-60 | (4) violin, piano, guitar, voice | (3) violin, piano and voice |
| 11 | 19 | 7 | 20 | 28 | (2) piano and voice | (1) voice |
| 12 | 51 | 3 | 12 | 65 | (3) piano, guitar and percussion | (1) guitar |
| 13 | 7 | 18 | 18 | 12 | (3) violin, piano and flute | (1) violin and piano |
| 14 | 30 | 8 | 28 | 35-56 | (1) piano | (1) piano |
| 15 | 28 | 14 | 6 | 21-42 | (3) voice, piano and guitar | (1) voice |
| 16 | 2 | 29 | 3 | 10 | (1) piano | (1) piano |

**Figure S.1.**
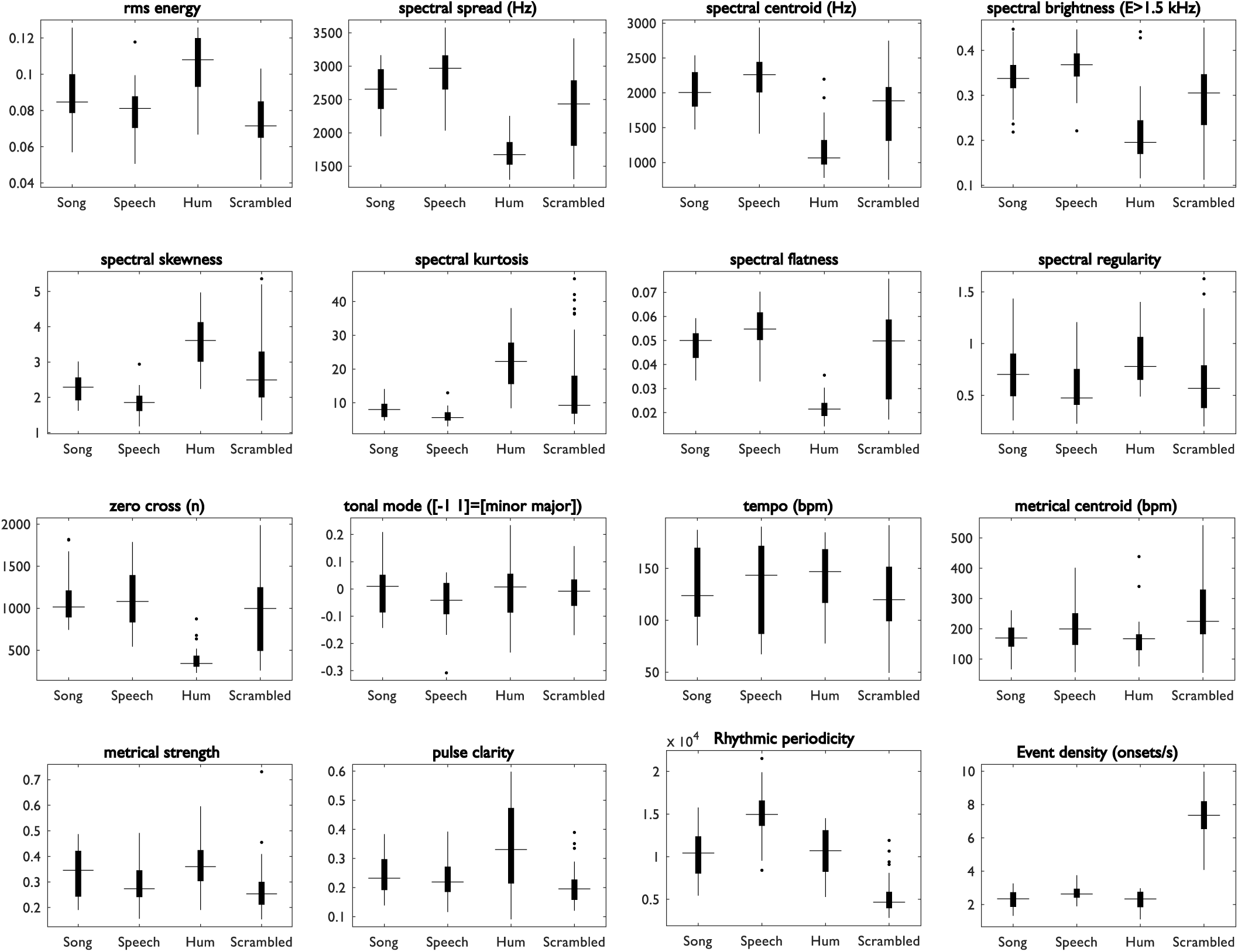
Box plot of acoustic characteristics across categories. Horizontal bars represent the median; black boxes show the interquartile range; vertical lines show the data range excluding outliers (dots).

**Figure S.2.**
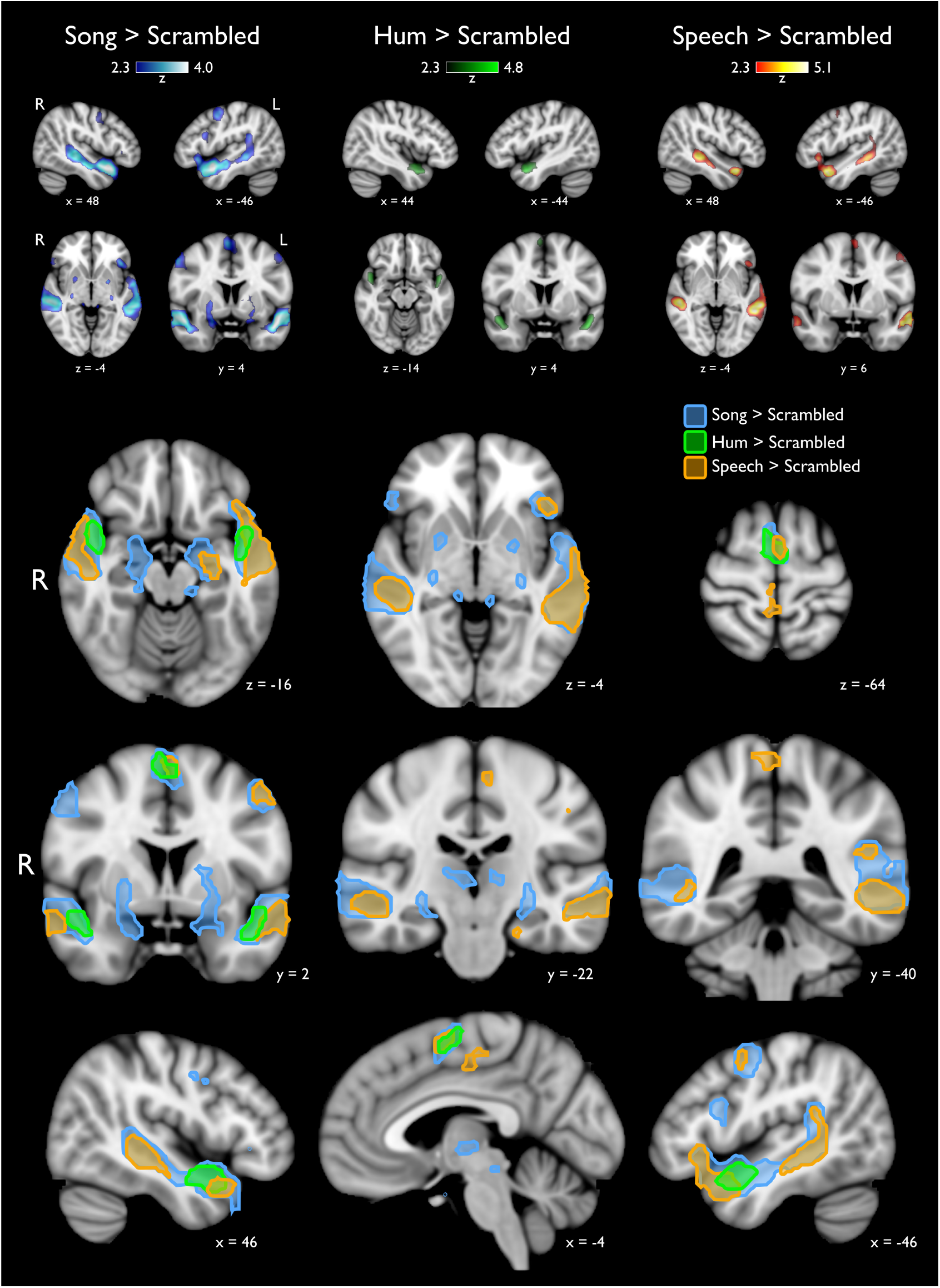
Natural stimuli versus overall Scrambled category (i.e., scrambled stimuli derived from the three original categories). Top panel is organized as in Figure 2. Bottom panel shows the outline of clusters resulting from the three contrasts indicated in the legend, thresholded at z>2.3, p_*cluster*_=0.05. Statistical maps are overlaid on the MNI-152 atlas. MNI coordinates of each slice are expressed in mm

## References

1. Fitch, W. T. The biology and evolution of music: A comparative perspective. Cognition 100, 173–215, doi:10.1016/j.cognition.2005.11.009 (2006).

2. MacLarnon, A. M. & Hewitt, G. P. The evolution of human speech: The role of enhanced breathing control. Am. J. Phys. Anthropol. 109, 341–363, doi:10.1002/(SICI)1096-8644(199907)109:3<341::AID-AJPA5>3.0.CO;2-2 (1999).

3. Patel, A. D. Why would Musical Training Benefit the Neural Encoding of Speech? The OPERA Hypothesis. Front. Psychol. 2, 142, doi:10.3389/fpsyg.2011.00142 (2011).

4. Patel, A. D. Can nonlinguistic musical training change the way the brain processes speech? The expanded OPERA hypothesis. Hear. Res. 308, 98–108, doi:10.1016/j.heares.2013.08.011 (2014).

5. Peretz, I., Vuvan, D., Lagrois, M.-E. & Armony, J. L. Neural overlap in processing music and speech. Philos. Transactions Royal Soc. London. Ser. B, Biol. Sci. 370, 20140090, doi:10.1098/rstb.2014.0090 (2015).

6. Pinker, S. How the Mind Works (W.W. Norton, 1999).

7. Koelsch, S. et al. Bach speaks: A cortical “language-network” serves the processing of music. NeuroImage 17, 956–966 (2002).

8. Merker, B. Synchronous chorusing and human origins. In The Origins of Music, 315–327 (The MIT Press, Cambridge, MA, US, 2000).

9. Schmithorst, V. J. Separate cortical networks involved in music perception: Preliminary functional MRI evidence for modularity of music processing. NeuroImage 25, 444–451, doi:10.1016/j.neuroimage.2004.12.006 (2005).

10. Schön, D. et al. Similar cerebral networks in language, music and song perception. NeuroImage 51, 450–461, doi:10.1016/j.neuroimage.2010.02.023 (2010).

11. Schön, D., Magne, C. & Besson, M. The music of speech: Music training facilitates pitch processing in both music and language. Psychophysiology 41, 341–349, doi:10.1111/1469-8986.00172.x (2004).

12. Zatorre, R. J. & Gandour, J. T. Neural specializations for speech and pitch: Moving beyond the dichotomies. Philos. Transactions Royal Soc. London. Ser. B, Biol. Sci. 363, 1087–1104, doi:10.1098/rstb.2007.2161 (2008).

13. Fitch, W. T. Four principles of bio-musicology. Philos. Transactions Royal Soc. London. Ser. B, Biol. Sci. 370, 20140091, doi:10.1098/rstb.2014.0091 (2015).

14. Fitch, W. T. Dance, Music, Meter and Groove: A Forgotten Partnership. Front. Hum. Neurosci. 10, 64, doi:10.3389/ fnhum.2016.00064 (2016).

15. Honing, H., ten Cate, C., Peretz, I. & Trehub, S. E. Without it no music: Cognition, biology and evolution of musicality. Philos. Transactions Royal Soc. London. Ser. B, Biol. Sci. 370, 20140088, doi:10.1098/rstb.2014.0088 (2015).

16. Bispham, J. C. Music’s “design features”: Musical motivation, musical pulse, and musical pitch. Music. Sci. 13, 41–61, doi:10.1177/1029864909013002041 (2009).

17. Tillmann, B. Pitch Processing in Music and Speech. Acoust. Aust. 42, 124–130 (2014).

18. Aichert, I., Späth, M. & Ziegler, W. The role of metrical information in apraxia of speech. Perceptual and acoustic analyses of word stress. Neuropsychologia 82, 171–178, doi:10.1016/j.neuropsychologia.2016.01.009 (2016).

19. Angulo-Perkins, A. et al. Music listening engages specific cortical regions within the temporal lobes: Differences between musicians and non-musicians. Cortex; a J. Devoted to Study Nerv. Syst. Behav. 59, 126–137, doi:10.1016/j.cortex.2014. 07.013 (2014).

20. Norman-Haignere, S., Kanwisher, N. G. & McDermott, J. H. Distinct Cortical Pathways for Music and Speech Revealed by Hypothesis-Free Voxel Decomposition. Neuron 88, 1281–1296, doi:10.1016/j.neuron.2015.11.035 (2015).

21. Rogalsky, C., Rong, F., Saberi, K. & Hickok, G. Functional anatomy of language and music perception: Temporal and structural factors investigated using functional magnetic resonance imaging. The J. Neurosci. The Off. J. Soc. for Neurosci. 31, 3843–3852, doi:10.1523/JNEUROSCI.4515-10.2011 (2011).

22. Leaver, A. M. & Rauschecker, J. P. Cortical representation of natural complex sounds: Effects of acoustic features and auditory object category. The J. Neurosci. The Off. J. Soc. for Neurosci. 30, 7604–7612, doi:10.1523/JNEUROSCI. 0296-10.2010 (2010).

23. Abrams, D. A. et al. Decoding temporal structure in music and speech relies on shared brain resources but elicits different fine-scale spatial patterns. Cereb. Cortex (New York, N.Y.: 1991) 21, 1507–1518, doi:10.1093/cercor/bhq198 (2011).

24. Merrill, J. et al. Perception of words and pitch patterns in song and speech. Front. Psychol. 3, 76, doi:10.3389/fpsyg. 2012.00076 (2012).

25. d’Errico, F. et al. Archaeological Evidence for the Emergence of Language, Symbolism, and Music–An Alternative Multidisciplinary Perspective. J. World Prehistory 17, 1–70, doi:10.1023/A:1023980201043 (2003).

26. De Souza, J. Voice and Instrument at the Origins of Music. Acad. Commons 97, 21–36, doi:10.7916/D8D50Q05 (2014).

27. Molino, J. Toward an evolutionary theory of music and language. In The Origins of Music, 165–176 (The MIT Press, Cambridge, MA, US, 2000).

28. Belin, P., Zatorre, R. J. & Ahad, P. Human temporal-lobe response to vocal sounds. Brain Res. Cogn. Brain Res. 13, 17–26 (2002).

29. Belin, P., Zatorre, R. J., Lafaille, P., Ahad, P. & Pike, B. Voice-selective areas in human auditory cortex. Nature 403, 309–312, doi:10.1038/35002078 (2000).

30. Callan, D. E. et al. Song and speech: Brain regions involved with perception and covert production. NeuroImage 31, 1327–1342, doi:10.1016/j.neuroimage.2006.01.036 (2006).

31. Ozdemir, E., Norton, A. & Schlaug, G. Shared and distinct neural correlates of singing and speaking. NeuroImage 33, 628–635, doi:10.1016/j.neuroimage.2006.07.013 (2006).

32. Brown, S., Martinez, M. J., Hodges, D. A., Fox, P. T. & Parsons, L. M. The song system of the human brain. Brain Res. Cogn. Brain Res. 20, 363–375, doi:10.1016/j.cogbrainres.2004.03.016 (2004).

33. Pantev, C., Roberts, L. E., Schulz, M., Engelien, A. & Ross, B. Timbre-specific enhancement of auditory cortical representations in musicians. Neuroreport 12, 169–174 (2001).

34. Pantev, C. et al. Increased auditory cortical representation in musicians. Nature 392, 811–814, doi:10.1038/33918 (1998).

35. Shahin, A., Bosnyak, D. J., Trainor, L. J. & Roberts, L. E. Enhancement of neuroplastic P2 and N1c auditory evoked potentials in musicians. The J. Neurosci. The Off. J. Soc. for Neurosci. 23, 5545–5552 (2003).

36. Shahin, A. J., Roberts, L. E., Chau, W., Trainor, L. J. & Miller, L. M. Music training leads to the development of timbre-specific gamma band activity. NeuroImage 41, 113–122, doi:10.1016/j.neuroimage.2008.01.067 (2008).

37. Fauvel, B. et al. Morphological brain plasticity induced by musical expertise is accompanied by modulation of functional connectivity at rest. NeuroImage 90, 179–188, doi:10.1016/j.neuroimage.2013.12.065 (2014).

38. Grahn, J. A. & Rowe, J. B. Feeling the beat: Premotor and striatal interactions in musicians and nonmusicians during beat perception. The J. Neurosci. The Off. J. Soc. for Neurosci. 29, 7540–7548, doi:10.1523/JNEUROSCI.2018-08.2009 (2009).

39. Strait, D. L., Chan, K., Ashley, R. & Kraus, N. Specialization among the specialized: Auditory brainstem function is tuned in to timbre. Cortex 48, 360–362, doi:10.1016/j.cortex.2011.03.015 (2012).

40. Herholz, S. C. & Zatorre, R. J. Musical Training as a Framework for Brain Plasticity: Behavior, Function, and Structure. Neuron 76, 486–502, doi:10.1016/j.neuron.2012.10.011 (2012).

41. Levitin, D. J. & Menon, V. Musical structure is processed in “language” areas of the brain: A possible role for Brodmann Area 47 in temporal coherence. NeuroImage 20, 2142–2152 (2003).

42. Roskies, A. L., Fiez, J. A., Balota, D. A., Raichle, M. E. & Petersen, S. E. Task-dependent modulation of regions in the left inferior frontal cortex during semantic processing. J. Cogn. Neurosci. 13, 829–843, doi:10.1162/08989290152541485 (2001).

43. Morosan, P., Schleicher, A., Amunts, K. & Zilles, K. Multimodal architectonic mapping of human superior temporal gyrus. Anat. Embryol. 210, 401–406, doi:10.1007/s00429-005-0029-1 (2005).

44. Woods, D. L. et al. Functional properties of human auditory cortical fields. Front. Syst. Neurosci. 4, 155, doi:10.3389/ fnsys.2010.00155 (2010).

45. Da Costa, S. et al. Human primary auditory cortex follows the shape of Heschl’s gyrus. The J. Neurosci. The Off. J. Soc. for Neurosci. 31, 14067–14075, doi:10.1523/JNEUROSCI.2000-11.2011 (2011).

46. Humphries, C., Liebenthal, E. & Binder, J. R. Tonotopic organization of human auditory cortex. NeuroImage 50, 1202–1211, doi:10.1016/j.neuroimage.2010.01.046 (2010).

47. Nudds, M. What Are Auditory Objects? Rev. Philos. Psychol. 1, 105–122 (2010).

48. Skipper, J. I. Echoes of the spoken past: How auditory cortex hears context during speech perception. Philos. Transactions Royal Soc. London. Ser. B, Biol. Sci. 369, 20130297, doi:10.1098/rstb.2013.0297 (2014).

49. Bookheimer, S. Functional MRI of language: New approaches to understanding the cortical organization of semantic processing. Annu. Rev. Neurosci. 25, 151–188, doi:10.1146/annurev.neuro.25.112701.142946 (2002).

50. Binder, J. R. et al. Human temporal lobe activation by speech and nonspeech sounds. Cereb. Cortex (New York, N.Y.: 1991) 10, 512–528 (2000).

51. Griffiths, T. D. & Warren, J. D. The planum temporale as a computational hub. Trends Neurosci. 25, 348–353 (2002).

52. Watkins, K. & Paus, T. Modulation of motor excitability during speech perception: The role of Broca’s area. J. Cogn. Neurosci. 16, 978–987, doi:10.1162/0898929041502616 (2004 Jul-Aug).

53. Chen, J. L., Zatorre, R. J. & Penhune, V. B. Interactions between auditory and dorsal premotor cortex during synchronization to musical rhythms. NeuroImage 32, 1771–1781, doi:10.1016/j.neuroimage.2006.04.207 (2006).

54. Chen, J. L., Penhune, V. B. & Zatorre, R. J. Listening to musical rhythms recruits motor regions of the brain. Cereb. Cortex (New York, N.Y.: 1991) 18, 2844–2854, doi:10.1093/cercor/bhn042 (2008).

55. Grahn, J. A. & Brett, M. Rhythm and beat perception in motor areas of the brain. J. Cogn. Neurosci. 19, 893–906, doi:10.1162/jocn.2007.19.5.893 (2007).

56. Merchant, H. & Honing, H. Are non-human primates capable of rhythmic entrainment? Evidence for the gradual audiomotor evolution hypothesis. Front. Neurosci. 7, 274, doi:10.3389/fnins.2013.00274 (2013).

57. Manning, F. & Schutz, M. “Moving to the beat” improves timing perception. Psychon. Bull. & Rev. 20, 1133–1139, doi:10.3758/s13423-013-0439-7 (2013).

58. Vuust, P. & Witek, M. A. G. Rhythmic complexity and predictive coding: A novel approach to modeling rhythm and meter perception in music. Front. Psychol. 5, 1111, doi:10.3389/fpsyg.2014.01111 (2014).

59. Friederici, A. D. & Alter, K. Lateralization of auditory language functions: A dynamic dual pathway model. Brain Lang. 89, 267–276, doi:10.1016/S0093-934X(03)00351-1 (2004).

60. Koelsch, S. & Siebel, W. A. Towards a neural basis of music perception. Trends Cogn. Sci. 9, 578–584, doi:10.1016/j. tics.2005.10.001 (2005).

61. Sammler, D., Grosbras, M.-H., Anwander, A., Bestelmeyer, P. E. G. & Belin, P. Dorsal and Ventral Pathways for Prosody. Curr. biology: CB 25, 3079–3085, doi:10.1016/j.cub.2015.10.009 (2015).

62. Price, C. J. The anatomy of language: A review of 100 fMRI studies published in 2009. Annals New York Acad. Sci. 1191, 62–88, doi:10.1111/j.1749-6632.2010.05444.x (2010).

63. Price, C. J. A review and synthesis of the first 20 years of PET and fMRI studies of heard speech, spoken language and reading. NeuroImage 62, 816–847, doi:10.1016/j.neuroimage.2012.04.062 (2012).

64. Ardila, A., Bernal, B. & Rosselli, M. How Localized are Language Brain Areas? A Review of Brodmann Areas Involvement in Oral Language. Arch. Clin. Neuropsychol. The Off. J. Natl. Acad. Neuropsychol. 31, 112–122, doi:10.1093/arclin/acv081 (2016).

65. Loucks, T. M. J., Poletto, C. J., Simonyan, K., Reynolds, C. L. & Ludlow, C. L. Human brain activation during phonation and exhalation: Common volitional control for two upper airway functions. NeuroImage 36, 131–143, doi:10.1016/j.neuroimage.2007.01.049 (2007).

66. Brown, S. et al. The somatotopy of speech: Phonation and articulation in the human motor cortex. Brain Cogn. 70, 31–41, doi:10.1016/j.bandc.2008.12.006 (2009).

67. Whitehead, J. C. & Armony, J. L. Singing in the brain: Neural representation of music and voice as revealed by fmri. Hum. Brain Mapp. 0, DOI: 10.1002/hbm.24333. https://onlinelibrary.wiley.com/doi/pdf/10.1002/hbm.24333.

68. Schneider, S., Schönle, P. W., Altenmüller, E. & Münte, T. F. Using musical instruments to improve motor skill recovery following a stroke. J. Neurol. 254, 1339–1346, doi:10.1007/s00415-006-0523-2 (2007).

69. Rodriguez-Fornells, A. et al. The involvement of audio-motor coupling in the music-supported therapy applied to stroke patients. Annals New York Acad. Sci. 1252, 282–293, doi:10.1111/j.1749-6632.2011.06425.x (2012).

70. Amengual, J. L. et al. Sensorimotor plasticity after music-supported therapy in chronic stroke patients revealed by transcranial magnetic stimulation. PloS One 8, e61883, doi:10.1371/journal.pone.0061883 (2013).

71. Chong, H. J., Cho, S.-R. & Kim, S. J. Hand rehabilitation using MIDI keyboard playing in adolescents with brain damage: A preliminary study. NeuroRehabilitation 34, 147–155, doi:10.3233/NRE-131026 (2014).

72. Gage, N. M. et al. Rightward hemispheric asymmetries in auditory language cortex in children with autistic disorder: An MRI investigation. J. Neurodev. Disord. 1, 205–214, doi:10.1007/s11689-009-9010-2 (2009).

73. Lai, G., Pantazatos, S. P., Schneider, H. & Hirsch, J. Neural systems for speech and song in autism. Brain: A J. Neurol. 135, 961–975, doi:10.1093/brain/awr335 (2012).

74. Paul, A. et al. The effect of sung speech on socio-communicative responsiveness in children with autism spectrum disorders. Front. Hum. Neurosci. 9, 555, doi:10.3389/fnhum.2015.00555 (2015).

75. Lartillot, O. & Toiviainen, P. MIR in Matlab (II): A toolbox for musical feature extraction from audio (Vienna, 2007).

76. Aubé, W., Angulo-Perkins, A., Peretz, I., Concha, L. & Armony, J. L. Fear across the senses: Brain responses to music, vocalizations and facial expressions. Soc. Cogn. Affect. Neurosci. 10, 399–407, doi:10.1093/scan/nsu067 (2015).

77. Beckmann, C. F., Jenkinson, M. & Smith, S. M. General multilevel linear modeling for group analysis in FMRI. NeuroImage 20, 1052–1063, doi:10.1016/S1053-8119(03)00435-X (2003).

78. Woolrich, M. W., Behrens, T. E. J., Beckmann, C. F., Jenkinson, M. & Smith, S. M. Multilevel linear modelling for FMRI group analysis using Bayesian inference. NeuroImage 21, 1732–1747, doi:10.1016/j.neuroimage.2003.12.023 (2004).

79. Worsley, K. J., Andermann, M., Koulis, T., MacDonald, D. & Evans, A. C. Detecting changes in nonisotropic images. Hum. Brain Mapp. 8, 98–101 (1999).

